# Targeted Gene Bisulfite Sequencing Identifies Differential Methylation in *p21*

**DOI:** 10.1101/291732

**Authors:** Ramya T. Kolli, Travis C. Glenn, Bradley T. Brown, Lillie M. Barnett, Lawrence H. Lash, Brian S. Cummings

## Abstract

Next-generation sequencing (NGS) methods are widely available to assess methylation of whole-genomes, reduced representation of genomes, and target capture of many loci, but simple, flexible, and low-cost methods are needed to leverage NGS for sequencing single-locus amplicons from large numbers of samples. We developed a two-stage PCR approach, targeted gene bisulfite sequencing (TGBS) which uses the Illumina MiSeq and Bismark bisulfite mapper, to assess site specific changes in methylation of the cyclin-dependent kinase inhibitor *p21* (*CDKN1a*) after exposure to a DNA methyltransferase inhibitor, 5-aza-2’-deoxycytidine (5-Aza) and determine the differences between human and rat *p21* methylation. TGBS analysis of human embryonic kidney cells (HEK293) and human proximal tubular cells (hPT) demonstrated variation at a known methylation sensitive site (SIE-1), but not in rat kidney cells. Treatment of cells with 5-Aza altered the methylation of this site in correlation with increased p21 protein expression. We also found that human and rat *p21* promoter sequences differ considerably in the amount of basal DNA methylation. These data showed the utility of TGBS for rapid analysis of DNA methylation of specific loci. We provide links to a ready-to-run Virtualbox that includes the program and commands for methylation analysis of bisulfite datasets, including step-by-step directions.

## Introduction

Next-generation sequencing (NGS) methods are widely used because they reduce the cost and time of data acquisition, enabling approaches such as whole-genome bisulfite sequencing (WGBS)^1^. The Illumina platform provides major advantages over the other existing technologies for DNA sequencing in general^2^ and is widely used for WGBS, reduced representation methylation sequencing^3^, and a variety of methods that target large numbers of loci (Supplementary Table S1). Single-locus DNA methylation analysis has traditionally been performed using cloning and Sanger sequencing^4,5^, but this method is labor intensive, time-consuming, and yields relatively low amounts of data^6^. NGS has replaced most applications that require large-scale cloning^2^. However, because most NGS methods have a high buy-in cost^7^, and most single-locus methylation analyses do not need huge amounts of data, the traditional cloning and Sanger bisulfite sequencing method is still commonly used.

Herein, we explain the work flow, kits, and optimized conditions suitable for the Illumina MiSeq and Bismark bisulfite mapping program to provide researchers with an all-in-one place user-friendly guide to targeted bisulfite sequencing. We discuss free software, such as Bismark, and focus on drastically reducing the time and cost of analysis. We also discuss Sanger sequencing and compared the data with NGS output. Our goal is to introduce the readers, especially those without NGS and/or bioinformatics support or experience, to the required instructions for affordable bench techniques and easy to reproduce sequence analysis tools, so that they can achieve cost- and time-efficient analysis.

Model System: DNA methylation adds methyl groups to the 5-carbon position of cytosine residues in the CpG dinucleotide context in multicellular eukaryotes. Being an epigenetic mark, the process doesn’t alter the core DNA sequence; however, depending on the location, it can regulate gene expression^8^. In general, promoter hypermethylation downregulates or silences gene expression^9^. Aberrant DNA methylation patterns in many genes have also been identified and correlated to the phenotypes of many cancers^10,11,12,13^. One such gene is *p21* (*CDKN1a*), a cyclin-dependent kinase (CDK) inhibitor observed to be silenced in many cancer by aberrant DNA methylation, as seen in metastatic prostate cancer, lung cancer and lymphomas^14,11,15,16^.

Large –omic based approaches yield excellent data on DNA methylation, but tend to be hypothesis generating studies describing hundreds, if not thousands of genes. Our previous research used omic-based approaches to identify *p21* as target gene for renal protection and the data suggested a role for DNA methylation in regulation of *p21*^17^. Thus, we needed a high-throughput technique to analyze DNA methylation at the *p21* promoter after exposure to various toxicants. A lack of validated high-through-put, rapid and robust approaches to assess gene targeted DNA methylation prompted us to develop the targeted gene bisulfite sequencing (TGBS) methodology. We used 5-Aza treatment because it has been previously shown to decrease methylation of *p21* promoter^10,18,19^ when assessed with traditional assays. We also used TGBS to investigate differences in DNA methylation between the promoter regions of human and rat *p21* genes.

## Methods

### Materials

Human embryonic kidney (HEK293) and normal rat kidney (NRK) cells and penicillin and streptomycin were purchased from American Type Culture Collection (Manassas, VA). Human proximal tubular (hPT) cells were derived from whole, de-identified human kidneys that were obtained through the International Institute for the Advancement of Medicine (Edison, NJ, USA). All tissue was scored by a pathologist as normal (i.e., derived from non-cancerous, non-diseased tissue). 5-aza-2’-deoxycytidine (5-Aza) and trypsin EDTA were purchased from Sigma-Aldrich (St. Louis, MO), DMEM media from HyClone technologies (Logan, UT), 5-Aza was dissolved in dimethyl sulfoxide (DMSO) from Fisher Scientific (Pittsburg, PA), DNeasy blood and tissue extraction kit were purchased from Qiagen (Valencia, CA). The EZ-DNA methylation lightning kit and the Zyppy plasmid miniprep kits were purchased from Zymo research (Irvine, CA). Nucleospin gel and PCR clean-up kits were purchased from Macherey-Nagel (Düren, Germany). The MiSeq reagent v3 kit was purchased from Illumina Inc (San Diego, CA), the Strataclone PCR cloning kit from Agilent technologies (Santa Clara, CA), the Kapa HiFi PCR kit from Kapa Biosystems (Wilmington, MA), and the Maxima hot-start Taq polymerase and Sera-Mag magnetic speedbeads were purchased from Thermo Scientific (Waltham, MA).

### Cell culture and treatment

5-Aza is a DNA methyltransferase inhibitor^20^ and is used in many studies for its demethylating properties^21,14,22^. HEK293 cells (3 × 10^6^) were seeded in T-175 tissue culture flasks and grown at 37°C in a 5% CO_2_ incubator for 24 hrs. Cells were then treated with 40 µM 5-Aza or DMSO (vehicle control for 5-Aza) for 72 hrs. The total amount of DMSO was never above 0.5% of the total volume per flask. Cells were released from the plate following treatment using trypsin/EDTA and 5×10^6^ cells were collected for DNA isolation.

hPT cell isolation procedure was based on that originally described by Todd et al.^23^, modified^24,25^, and reported in Huang et al.^26^. Briefly, sterile conditions (i.e., all instruments and glassware were autoclaved, and all buffers were filtered through a 0.2-μm pore-size filter) were used. Renal cortex and outer stripe were cut into slices, washed with sterile PBS, minced, and the pieces were placed in a trypsinization flask filled with 300 ml of sterile, filtered Hanks’ buffer, containing 25 mM NaHCO_3_, 25 mM HEPES, pH 7.4, 0.5 mM EGTA, 0.2% (w/v) bovine serum albumin, 50 μg/ml gentamicin, 1.3 mg/ml collagenase, and 0.59 mg/ml CaCl_2_, which was filtered prior to use. Whole kidneys were perfused with Wisconsin or similar type medium and kept on ice until they arrived at the laboratory, which was usually within 24 h of removal from the donor. All buffers were continuously bubbled with 95% O_2_ / 5% CO_2_ and were maintained at 37°C. Minced cortical pieces from whole kidneys were subjected to collagenase digestion for 60 min, after which the supernatant was filtered through a 70-μm mesh filter to remove tissue fragments, centrifuged at 150 × *g* for 7 min, and the pellet suspended in Dulbecco’s Modified Eagle’s Medium: Ham’s F-12 (DMEM/F-12; 1:1). Approximately 5 to 7 × 10^6^ cells were obtained per 1 g of human kidney cortical tissue.

### DNA extraction and bisulfite conversion

Cells (5×10^6^) were pelleted at 1000 rpm for 5 min and the supernatant was discarded. Genomic DNA was extracted using the Qiagen’s DNeasy blood and tissue kit following the manufacturer’s protocol. DNA was eluted in two successive steps to obtain a maximum yield, using 120 µl followed by 40 µl of elusion buffer. Following quantification using a Nanodrop spectrophotometer, 2 µg of the extracted DNA was bisulfite treated using the Zymo Research’s EZ-DNA methylation lightning kit following the manufacturer’s protocol, and re-quantified.

### Target amplification and purification

Bisulfite converted DNA (350 ng) was used to amplify different regions of the *p21* promoter. The locus specific primers were designed using Methprimer^27^ and were synthesized by Integrated DNA Technologies Inc. (IDT, Coralville, IA). Partial TruSeqHT sequences corresponding to part of the Illumina Read1 (R1) and Illumina Read2 (R2) sequencing primer-binding sites^28,29,30^ were added 5’ to the locus specific primers during primer synthesis. The locus specific primers and the partial TruSeqHT sequences are as given in **Table 1**. Fusion primers were synthesized by IDT where R1 was fused to forward primers and R2 was fused to reverse primers. For example, the primer pair for hp21-TSS site was, forward: “iTru R1 + hp21-TSS F” (ACACTCTTTCCCTACACGACGCTCTTCCGATCT ATAGTGTTGTGTTTTTTTGGAGAGTG) and reverse: “iTru R2 + hp21-TSS R” (GTGACTGGAGTTCAGACGTGTGCTCTTCCGATCTACAACTACTCACACCTCAACTAAC).

**Table 1.**
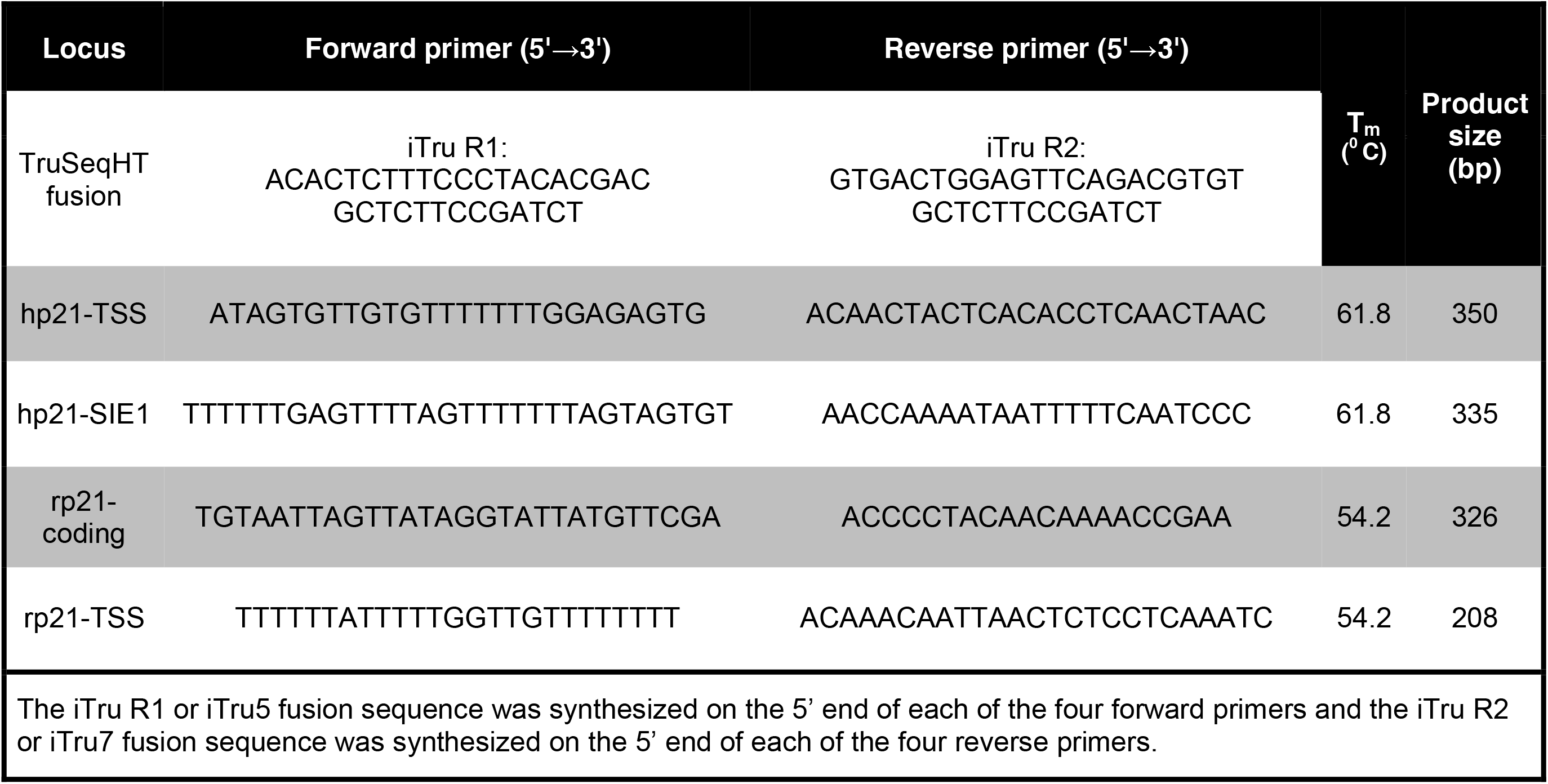
TruSeqHT fusion stubs and locus-specific primers.

The first locus amplified was a 350 bp fragment of the human *p21* promoter region adjacent to the transcription start site (TSS) termed as hp21-TSS. The second locus was a 335 bp fragment including the transcription factor binding site approximately 700 bp upstream of the TSS called the sis-inducible element (SIE-1) termed hp21-SIE1. The third site was the 208 bp fragment of the rat *p21* promoter region near the TSS termed as rp21-TSS, and the fourth locus was the 326 bp rat *p21* coding region approximately 9 Kb downstream of the TSS, termed rp21-coding (**Figure 1, Table 1**).

**Figure 1.**
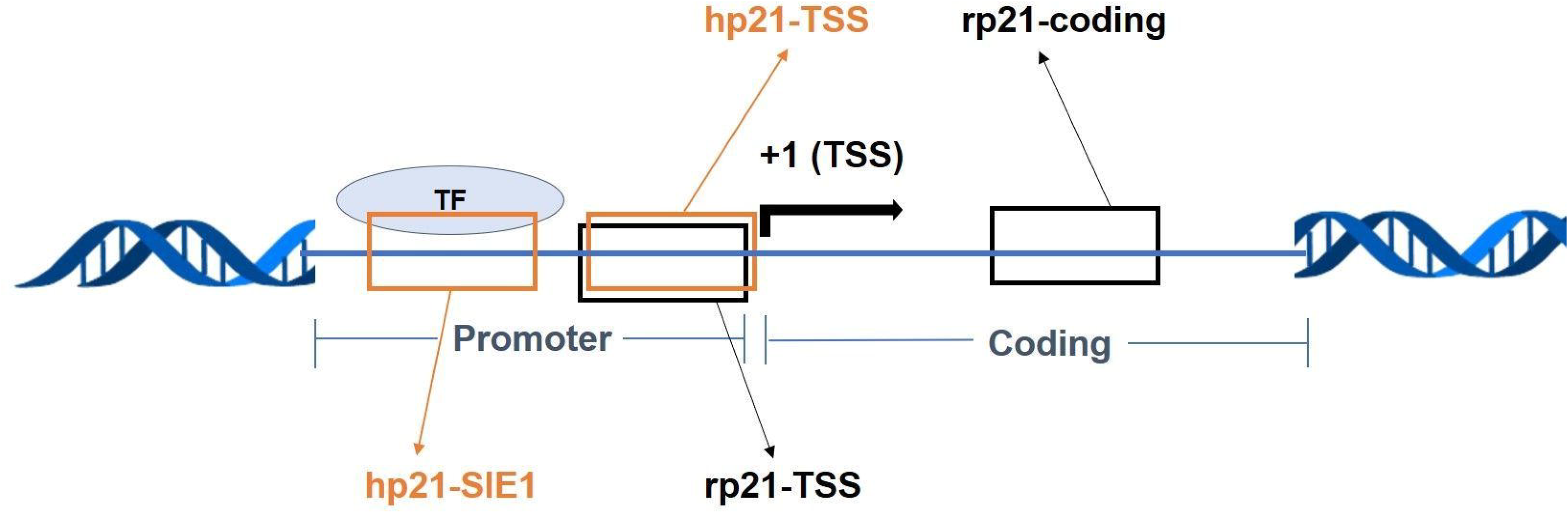
Schematic of *p21* gene organization highlighting the loci of interest for DNA methylation analysis. This includes the human *p21* promoter region adjacent to the transcription start site (hp21-TSS), the human transcription factor binding site called the sis-inducible element (hp21-SIE1), the rat *p21* promoter region starting near the start site (rp21-TSS) and the rat *p21* coding region (rp21-coding).

The 25 µl PCR amplification reaction mix contained 3 mM MgCl_2_, 1X hot start buffer (Thermo Scientific), 0.2 mM of each deoxynucleoside 5’-triphosphate (dNTP), 0.4 µM each of the forward and reverse primers, 1.5 units HotStart Taq DNA polymerase (Thermo Scientific) and the 350 ng DNA template. PCR was performed under the following conditions: 95°C for 5 min, 40 cycles of 95°C for 30 sec, 61.8°C (hp21-TSS, hp21-SIE1) or 54.2°C (rp21-TSS or rp21-coding) for 45 sec followed by, 72°C for 45 sec and a final 72°C for 10 min. The PCR products were then separated by electrophoresis on a 1% (w/v) agarose gel and visualized with ethidium bromide under a UV trans-illuminator and the amplicons corresponding to the loci were extracted from the gel using Nucleospin gel and PCR clean-up kit (Macherey-Nagel) following the manufacturer’s instructions. The sequences of the purified PCR products were confirmed using Sanger sequencing at the Georgia Genomics and Bioinformatics Core (GGBC) at the University of Georgia. All sequences obtained were verified for locus-specificity using the Basic Local Alignment Search Tool (BLAST)^31^.

### Sanger sequencing of bacterial clones

StrataClone PCR cloning kits (Agilent) were used for Sanger bisulfite sequencing. Briefly, 50 ng of gel extracted PCR products were cloned into *Escherichia coli* (*E. coli*) following the manufacturer’s instructions. After cloning and plating, about 3 bacterial colonies (white or light blue) were picked and suspension cultures were prepared for plasmid minipreps. Plasmids from the cultures were isolated using Zyppy plasmid miniprep kit (Zymo Research) following the manufacturer’s instructions. The plasmid inserts were sequenced at the GGBC by Sanger sequencing. The sequences were analyzed using BiQ Analyzer DNA methylation analysis software following software instructions^32^.

### Library preparation and next-generation sequencing

Purified PCR amplicons from agarose gel extraction were normalized to 5 ng/µl. A limited cycle PCR was performed to attach the iTru5 and iTru7 primers with eight nucleotide indexes, as represented in **Figure 2**. The 25 µl limited cycle reaction contained 1X Kapa buffer, 0.3 mM of each dNTP, 0.3 µM of each primer, 25 ng template DNA and 0.5 U of HiFi hotstart DNA polymerase (Kapa Biosciences). The reaction conditions were: 98°C for 5 min, 11 cycles of 98°C for 15 sec, 60°C for 30 sec and 72°C for 30 sec and a final 72°C for 1 min. Aliquots (10 µl) from each reaction were pooled together and cleaned up using Thermo Scientific’s Sera-Mag magnetic speedbeads. An equal ratio of speedbeads to the sample pool were vortexed and placed on the magnet and incubated at room temperature for 10 min. Once the beads were drawn to the magnet, the supernatant was discarded. The beads were washed with 80% ethyl alcohol twice and the residual liquid was removed by absorption using a wooden toothpick. DNA was then eluted using TLE buffer (10 mM Tris pH 8 & 0.1 mM EDTA) and supernatant collected. The pooled and cleaned sample was then processed for sequencing on an Illumina MiSeq platform as described by Glenn *et al*^29^ using the Illumina’s MiSeq 600 cycle v3 kit.

**Figure 2.**
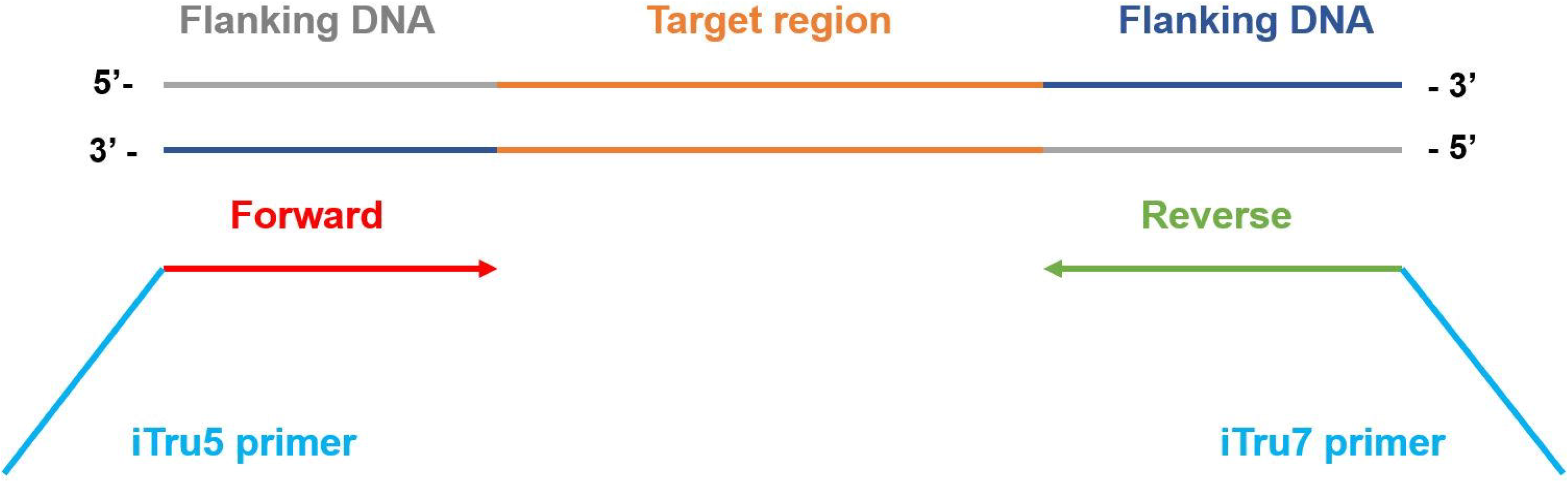
TruSeqHT fusion primers for Illumina MiSeq platform. The forward and reverse portions of the primers are the locus-specific primers that are complimentary to the flanking regions. iTru5 and iTru7 primers with unique index combinations allow identifying the source samples in the sequence pool.

## Sequence analysis

### Read quality and trimming

The paired end 250-350 bp reads obtained from Illumina MiSeq were demultiplexed using Illumina software bcl2fastq^33^. The sequence reads in fastq format were trimmed for better alignment using Babraham Bioinformatics’ free software Trim Galore^34^ or Geneious^35,36^. However, because the sequencing templates included mostly the uniformly sized PCR products, trimming did not affect read alignment (data not shown).

### DNA methylation analysis using Bismark

Bismark bisulfite mapper is a Linux based free software from Babraham Bioinformatics Institute^37^. Methylation analysis using Bismark was carried out in three steps:

#### 1. Genome preparation

A reference genome was prepared where an NCBI genome sequence for the target locus (*p21* promoter or coding region) was downloaded as a fasta file. The reference genome was prepared using the following command from the software guide: “/bismark/bismark_genome_preparation --path_to_bowtie /usr/bin/bowtie2/ --verbose /data/genomes/homo_sapiens/GRCh37/”. For example: “bismark_genome_preparation --/home/user/DNA/bowtie2-2.3.0/ --verbose /home/user/DNA/bowtie2-2.3.0/bismark_v0.17.0/REF/”, where the reference fasta file was saved in a directory or folder REF in the home folder of the user within the bismark folder. This created two folders within the genome folder REF, one with C ->T genome index and another with G ->A for the reverse reads.

#### 2. Read alignment

The second step was running bowtie2 within Bismark using the command: “bismark –bowtie2 -n 1 -l 50 /data/genomes/homo_sapiens/GRCh37/ test_dataset.fastq”. For example, read alignment for sequences in the folder named SampleSeq_R1 with a single-end approach was performed using: “bismark --bowtie2

/home/user/DNA/bowtie2-2.3.0/bismark_v0.17.0/REF/ SampleSeq_R1.fastq.gz”. This aligned the sequence reads to the reference genome and created a combined alignment or methylation call output in a binary representation of sequence alignment map called BAM format, and yielded a run statistics report. Output files included a bam file and report.txt file (**Supplementary Datafile 1**). The BAM file can only be opened in Bismark.

#### 3. Methylation extraction

The third and final step was methylation extraction of the bam file generated in the second step. The command used was “bismark_methylation_extractor --gzip test_dataset.fastq_bismark.bam”. For example: “bismark_methylation_extractor -s – comprehensive SampleSeq_R1.fastq_bismark_bt2.bam”. This generated output files that included M-bias.txt file, M-bias_R1.png file, CpG (**Supplementary Datafile 2-4**), CHH and CHG context bt2.txt files, which contain information on strand specific methylation. The key information on CpG site specific percent methylation was obtained from the M-bias.txt file (**Supplementary Datafile 2**). The targeted bisulfite sequencing with short products allowed for manual extraction of methylation values for comparison across samples and treatments. A processing report was generated using the command “bismark2report” that summarized the process with a read alignment chart, methylation extraction report and an M-bias plot.

### VirtualBox with ready-to-run Bismark package

VirtualBox is an open source software that runs on various operating systems and supports various guest operating systems^38^. The path to download a ready-to-run VirtualBox package containing all the tools and installations required for DNA methylation analysis of a given fastq sequence file is indicated below. The package includes step-by-step instructions for running Bismark, which is a Linux software, in a VirtualBox on Windows host system. This can be found at “http://toxicology.uga.edu/resources/dna_methylation_analysis/” for the VirtualMachine named “TGBS” that can be accessed with the username “user” and password “TGBSKolli”. The instructions are detailed in the **Supplementary File**.

#### Statistics

Samples isolated from a distinct cell passage represented one experiment (n=1). Data are represented as mean ± SEM (standard error of the mean) from at least three separate experiments (n=3). An unpaired Student’s t-test was used to compare two groups using Graphpad PRISM considering p<0.05 indicative of a statistically significant difference between the mean values.

## Results and Discussion

### Sanger vs next-generation bisulfite sequencing

Normal rat kidney (NRK) cells were treated with 40 µM 5-Aza as a positive control, or its vehicle control (DMSO) for 72 hrs as previously described^39^. The extracted DNA was subjected to bisulfite conversion and the rp21-coding region was amplified using the primers described in **Table 1**. The PCR products were separated by electrophoresis and then gel purified. The 350 bp purified products were cloned into competent bacteria for further target enrichment. Plasmids from three clones per treatment were individually sequenced by the Sanger tube method. The sequences obtained were either manually aligned or aligned using BiQ Analyzer software (**Figure 3A**). In general, at least one out of three clones showed a different methylation pattern leading to inconsistency and resulting in the need for more clones to reach an acceptable power of analysis, irrespective of the treatment. Typically, this would require an average of 8-10 clones per treatment group.

**Figure 3.**
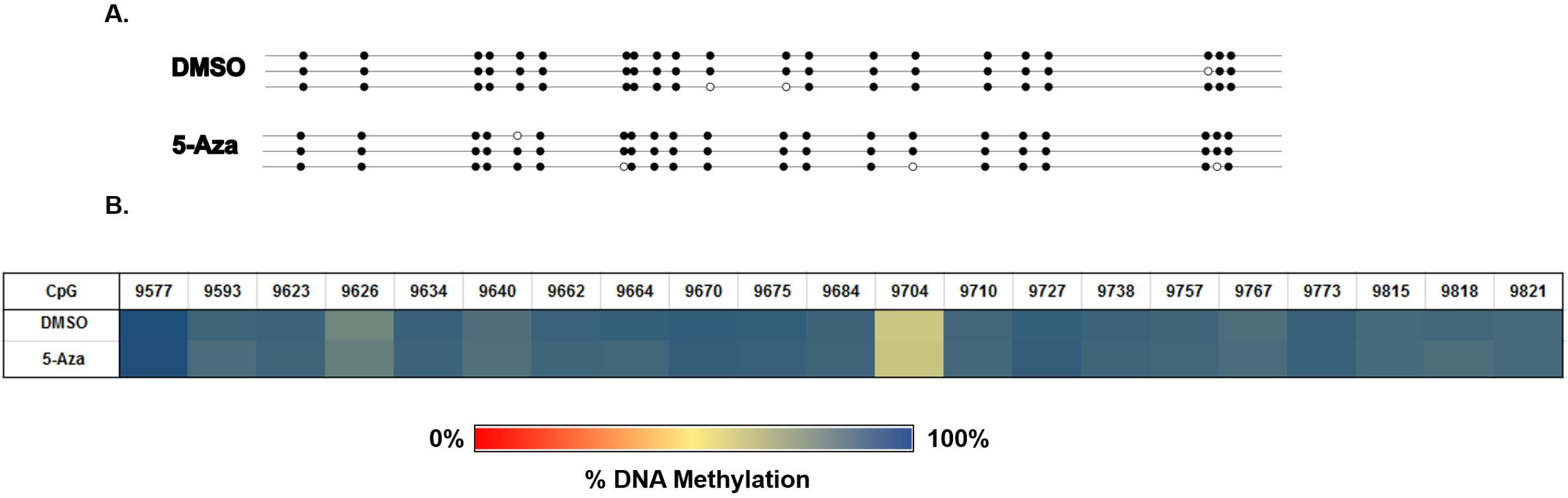
DNA methylation status of the rat *p21* coding region. **A)** Methylation of the rat *p21* coding region as determined using Sanger bisulfite sequencing: Data are shown as a lollipop plot generated using BiQ Analyzer. Each treated group includes three random clones and each line represents sequence from a clone. Black indicates methylated CpGs and the white represents unmethylated CpGs. **B)** Methylation of the rat *p21* coding region as determined by TGBS using Illumina next-generation sequencing: Data are represented as a heat-map with average DNA methylation increasing from red (0%) to blue (100%). The position indicates the CpG dinucleotide site in the sequenced fragment.

To address the variation among clones assayed by Sanger sequencing, we sequenced the same rp21-coding region on the Illumina MiSeq platform using TGBS. The gel purified 350 bp products were subjected to limited cycle PCR to attach the iTru5 and iTru7 primers with unique index combinations for each sample. All the samples were pooled, purified and then sequenced using the 600 cycle v3 kits. An average of about 10,000 reads were obtained per sample, hence increasing the statistical power of analysis^40^ as compared to the Sanger sequencing. The data from Bismark’s text file outputs were compiled together and the percent DNA methylation changes were displayed using heat-maps (**Figure 3B**). The methylation status of this coding region did not change after treatment with 5-Aza, suggesting that the coding region is not one of the key components in epigenetic regulation of *p21* expression. Nevertheless, in addition to being derived from about 10,000 reads per sample, the data showed consistency across three independent experiments.

## Differential methylation analysis

### Differences in basal DNA methylation of the *p21* promoter region between human and rat kidney cells

We used TGBS to assess differences in basal DNA methylation between human and rat *p21* promoters isolated from HEK293 and NRK cells. This included analysis of a 350 bp upstream fragment of the human *p21* promoter adjacent to the transcription start site (hp21-TSS) and a 250 bp upstream fragment of the rat *p21* promoter near the transcription start site (rp21-TSS). The data showed differences in methylation between the two cell lines with an average of 16.4% at the rp21-TSS site and 0.8% at the hp21-TSS site (**Figure 4A-B**), suggesting species-dependent differences in basal methylation of the *p21* promoter region at the TSS. This is a critical point as it further suggests that epigenetic changes in rat genes cannot be used as a surrogate for human genes, a key consideration in using epigenetics in rats to infer changes in human genes.

**Figure 4.**
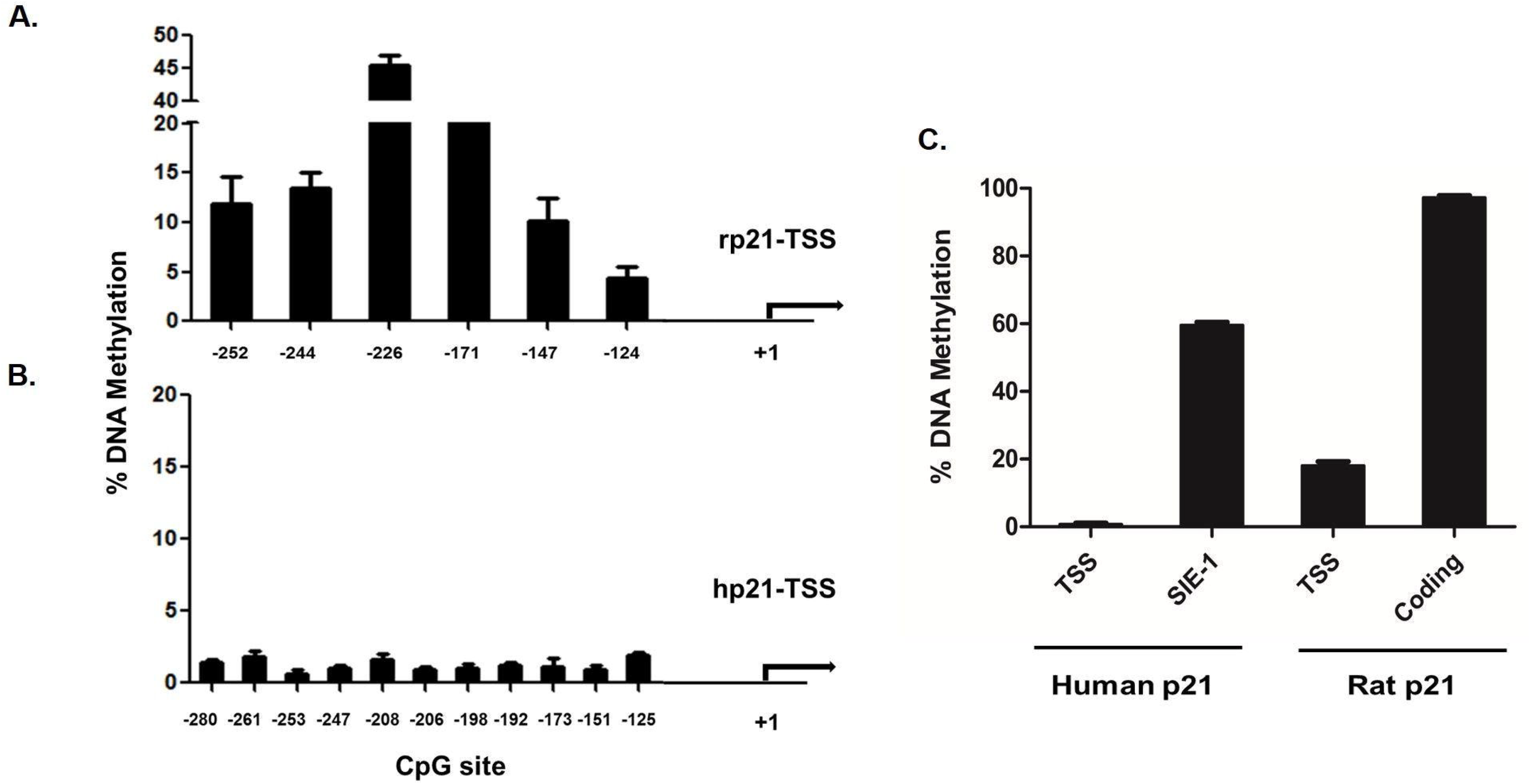
Differential methylation analysis. Comparison of methylation between rat (**A**) and human (**B**) *p21* transcription start sites. DNA methylation data are represented as percent methylation of each CpG site in the analyzed fragments of the human and rat *p21* promoter regions near the respective transcription start sites (rp21-TSS and hp21-TSS). (**C**) Comparison of methylation in different regions of the rat and human *p21* gene. Differential methylation data are represented as percent DNA methylation of the transcription start site (TSS), sis-inducible element (SIE-1) and gene coding regions of human and rat *p21*. Next-generation sequencing data was analyzed using Bismark bisulfite mapper. Data are represented as the mean ± standard error of the mean (SEM) of three independent experiments (n=3).

### Regional differences in basal DNA methylation of p21

A differential methylation analysis was performed on *p21* DNA isolated from untreated cells. The regions analyzed are as shown in **Figure 1**. Across the two species, we observed differential methylation across these regions with an average of 0.8% at the hp21-TSS, 57.9% at the transcription factor binding site called sis-inducible element (SIE-1), 16.1% at the rat-TSS and 95.8% at the rat p21-coding region (**Figure 4C**). This not only highlights the regional differences in DNA methylation of *p21*, but also validates and confirms the repeatability and reproducibility of TGBS for such analysis.

### Effect of 5-Aza

We used 5-Aza as a positive control to verify the ability of TGBS to detect changes in DNA methylation. The basal level of methylation in the CpG sites spanning 350 bp adjacent to the human *p21* transcription start site (hp21-TSS) showed a low level of total percent methylation (0.9%) in the presence of DMSO, which was similar to the 5-Aza treated cells (**Figure 5A**). In general, promoter regions near the TSSs are rich in CpG islands (GC% > 50), which are known to be typically unmethylated in normal cells for active transcription to persist^41^. The low level of methylation detected in the human *p21* promoter region most likely explains the relative infectiveness of 5-Aza, as a two-fold change would only result in decrease of 0.45%. To address this, we also assessed methylation upstream of the TSS at a transcription factor binding site about 700 bp from the start site CpG islands, called the sis-inducible element (SIE-1). This site is recognized by members of the signal transducer and activator of transcription family (STAT). The binding of STAT1 protein to SIE-1 has also been shown to upregulate *p21* expression^42^.

**Figure 5.**
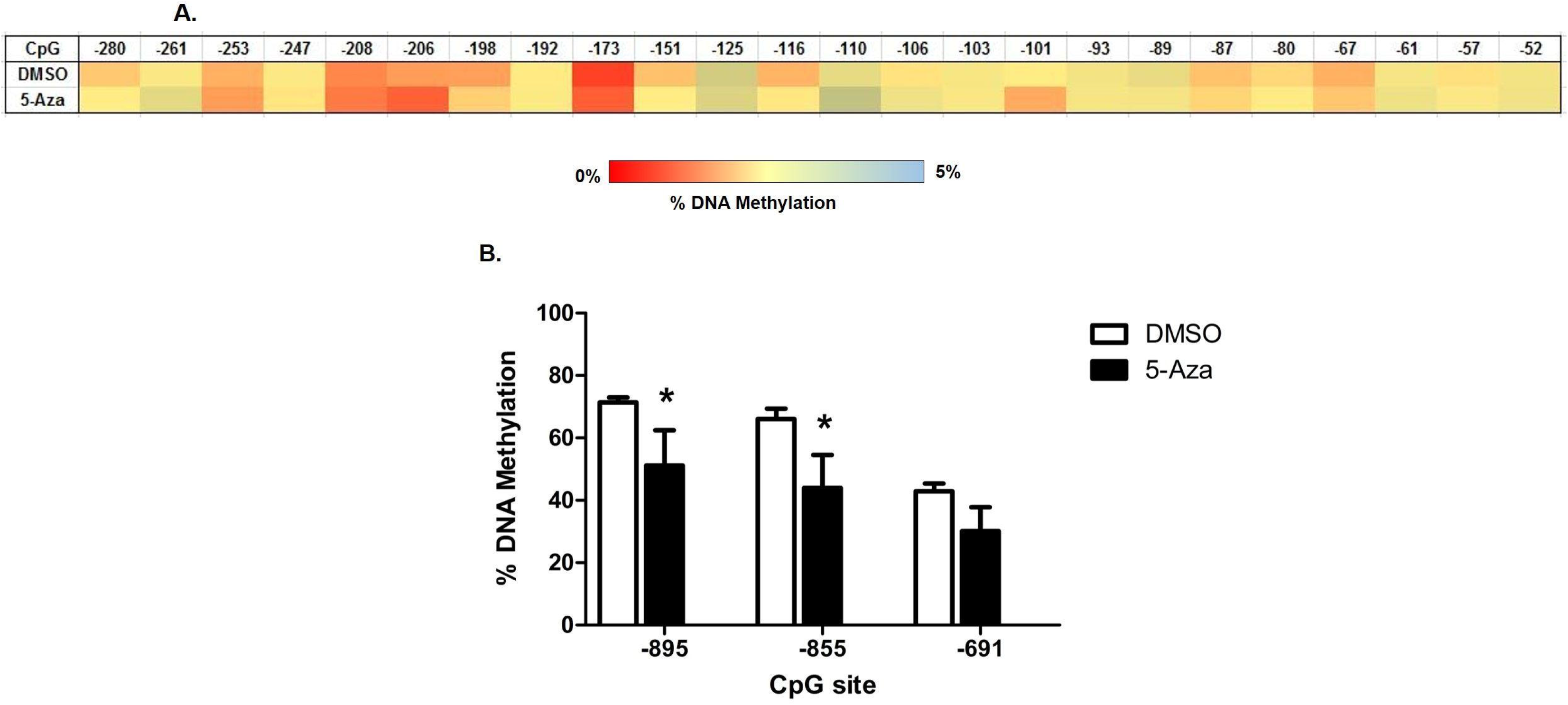
Effect of 5-Aza on DNA methylation of the promoter region of human *p21*. **(A)** Heat-map of the site-specific percent DNA methylation changes as determined by TGBS in the human *p21* promoter region at the transcription start site (hp21-TSS) after 3 days of exposure to DMSO (vehicle control) or 40 µM 5-Aza (positive control). The first row represents the position of the cytosine in the CpG dinucleotide context relative to the TSS. Heat map intensity is showed in the sidebar with deep red indicating percent methylation value towards zero and pale blue indicating relatively higher methylation of 5%. (**B)** Effect of 5-Aza on DNA methylation of cytosine residues in the SIE-1 site in human *p21* promoter region (hp21-SIE1). Data are represented as the mean ± SEM of three independent experiments (n=3). *P<0.05 compared with DMSO.

Treatment of cells with 5-Aza caused about a 35% decrease in total methylation of the SIE-1, as compared to DMSO treated cells (**Figure 5B**). This correlated to increases in the protein expression of p21 as shown in our previously published study^39^. This validates the site SIE-1 to be an important region for investigating the mechanism of action behind the epigenetic regulation of *p21* expression and more importantly validates our method developed.

### Differences in basal DNA methylation of the *p21* promoter region between HEK293 cells and freshly isolated human proximal tubule cells

As we observed decreases in the percent DNA methylation at the hp21-SIE1 site after treatment with the demethylating agent 5-Aza, we wanted to investigate the differences in basal level methylation between HEK293 cells and freshly isolated human proximal tubule (hPT) cells at the TSS and sites. The average methylation of hp21-TSS in hPT cells was 1.4% and not significantly different from that in HEK293 cells **(Figure 6A)**. In contrast, the average methylation of all three CpG sties in hPT cells were lower than that measured in HEK293 cells at the hp21-SIE1 site **(Figure 6B)**.

**Figure 6.**
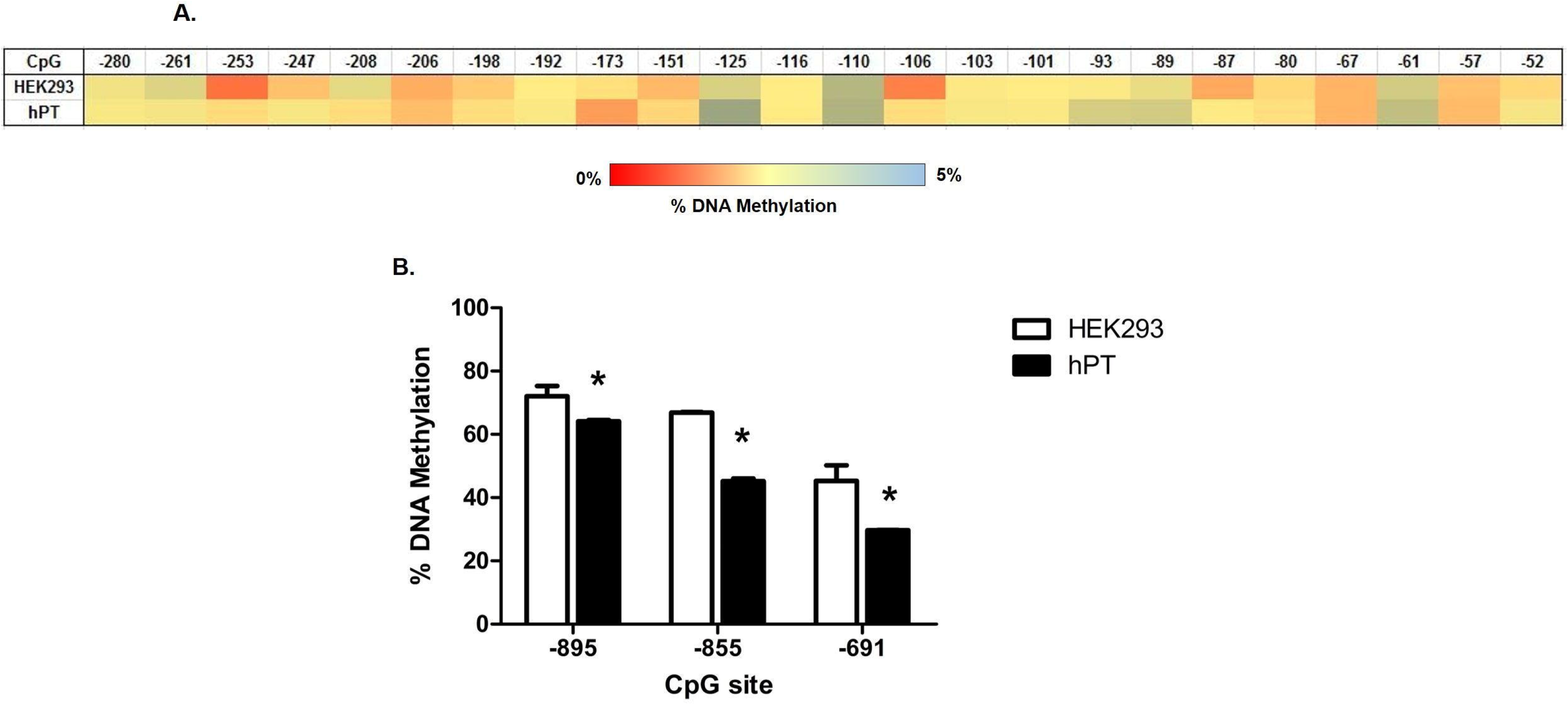
Comparison of basal DNA methylation of the *p21* promoter region between HEK293 cells and freshly isolated human proximal tubule (hPT) cells. **(A)** Heat-map of the site-specific percent DNA methylation changes as determined by TGBS in the human *p21* promoter region at the transcription start site (hp21-TSS). Heat map intensity is showed in the sidebar with deep red indicating percent methylation value towards zero and pale blue indicating relatively higher methylation of 5%. **(B)** Comparison of methylation of human *p21* promoter at the transcription factor binding site SIE-1 between HEK293 and hPT cells. Data are represented as the mean ± SEM of three different passages of HEK293 cells and three different pools of the hPT isolated cells (n=3). *P<0.05 compared with HEK293.

### Advantages of TGBS

Currently, over 90% of the world’s sequencing data is produced using Illumina technologies. Thus, it is expected that Illumina sequencing is beginning to see more application in the field of epigenetics. As such, various Illumina instruments are used to produce bisulfite DNA methylation data including MiSeq, HiSeq and NextSeq instruments. WGBS requires high numbers of reads per sample (e.g., ≥1 million reads), but because these are distributed across the genome, the read depth per nucleotide is relatively low (e.g., ≤50 x). In contrast, TGBS uses relatively low number of sequencing reads per sample (i.e., ≤10,000 reads), but because the reads are targeted to the locus of interest, read depths per nucleotide are quite deep (e.g., ≥1000x). The limited number of reads per sample allowed for rapid data analysis, far lower costs per sample, but higher precision in estimating methylation.

The use of TGBS, as opposed to Sanger or pyrosequencing, provides major advantages over the other existing technologies in general^2^. Recent studies, including our own^39^ have used methylation-specific PCR to analyze changes in DNA methylation; allowing us to only assess change in methylation of two CpG sites within the primer binding region. In contrast, TGBS facilitated assessing ~10,000 reads of each *p21* locus from hundreds of samples in a single run, at costs similar to cloning and sequencing a few copies of each gene from a few samples using traditional cloning and Sanger sequencing. The differences in the work flow between first-generation Sanger bisulfite sequencing and next-generation TGBS is described in **Supplementary Figure S1**. By combining the TGBS approach with standard statistical approaches (multiple passages and samples) a more robust approach is evident.

In summary, we combined two different existing technologies to develop a novel approach to rapidly identify specific CpG sites whose methylation is altered in the nephron-protective gene *p21*. The positive results with 5-Aza suggest that the SIE-1 is an important region for epigenetic regulation of *p21* expression. This assertion is strengthened by the fact that we have already shown that 5-Aza alone induced p21 protein expression in these cells^43^. To our knowledge this is the first report demonstrating the methylation of specific DNA bases within *p21* promoter that are altered by 5-Aza exposure. We also used this technique to demonstrate differences in the basal promoter methylation between rat and human *p21*. These data also allowed us to compare HEK293 cells to freshly isolated human proximal tubule cells. A significant difference in basal DNA methylation was observed between the two cell types. However, the implication of these difference in terms of cellular function or response to pathological stimuli, or toxicant exposure has yet to be investigated.

## Acknowledgements

We thank everyone involved in the generation of data. We thank Troy J. Kieran and Patrick R. Watson (T.C.G. laboratory) for their help in sample pooling and MiSeq runs. This project was supported in part with funds from the National Institutes of Health (NIH); National Institute of Biomedical Imaging and Bioengineering (NIBIB (EB0160100 to B.S.C.), as well as Interdisciplinary Toxicology Program Stipend Support and US - EPA’s Student Services Authority Support to R.T.K.

