## Supplementary Materials for "Targeted Gene Bisulfite Sequencing Identifies Differential Methylation in *p21*"

### Supplementary File

INSTRUCTIONS FOR RUNNING BISMARK IN VIRTUALBOX

#### 1. System requirements

- A minimum of 2GB RAM for VirtualBox guest operating system with a preferred host system total of 8GB RAM or greater.
- A minimum of 10GB disk storage.
- “Virtualization Technology” is supported in the host system BIOS.  Set it to Enable.

#### 2. VirtualBox installation

- Download the latest VirtualBox version for your host operating system:

  https://www.virtualbox.org/wiki/Downloads

Follow the instructions and install VirtualBox.

#### 3. Download image file containing Bismark tool-set

- Open any web browser on the host system and go to: http://toxicology.uga.edu/resources/dna_methylation_analysis/
- Download the “TGBS.ova” file and save it on the host system.
- In the VirtualBox Manager, go to “File” and “Import Appliance” and browse for the location of the “TGBS.ova” file.

#### 3. Run the VirtualMachine

- Start the “TGBS machine”. If error occurs, go to “Settings”, select “USB” and disable the “USB controller”. Then, start the “TGBS machine” again.
- Use password **TGBSKolli** for the **User** login.

- Click on the “Activities” tab and open “Files”.
- The home directory includes the Bismark software version 0.17.0 and the required tools like ActivePearl. It also includes a sample fastq sequence file “SampleSeq_R1.fastq.gz”.

**4. Download Reference Genome:**

- Select “Firefox” from the “Activities” tab and go to https://www.ncbi.nlm.nih.gov/
- In the search bar, select “Nucleotide” from the drop-down menu and then search for “U24170.1”
- Click on “FASTA” and select the option to “Send” the “Complete Record” to the “File” destination and select “Create File” and “Save File”.
- The “Sequence FASTA” is now in the “Downloads” folder.
- Using the Files tool again, move sequence.fasta to:

    Home->DNA->bowtie2-2.3.0->bismark_v0.17.0->REF->.

**5. DNA Methylation analysis**

Following are the command lines for a single-end analysis using R1 forward reads.

- Note: Click on the “Activities” tab to move between the folders.

**5.1. Prepare Reference Genome:**

- Select “Terminal” from the “Activities” tab.
- The reference genome folder “REF” now contains a fasta file from NCBI with the Accession# U24170.1 for the human *p21* promoter. Use the following command to prepare the reference genome before mapping the bisulfite converted reads

from the sample datafile. Note: In the command below there is a single space before each "--" (double hyphen) and a single space after verbose.

```
bismark_genome_preparation --/home/user/DNA/bowtie2-2.3.0/ --verbose
/home/user/DNA/bowtie2-2.3.0/bismark_v0.17.0/REF/
```

- This creates two folders within the genome folder "REF", one with C ->T genome index and another with G ->A.

**5.2. Run Bismark:**

- Read alignment step for sequences in the R1 read file "SampleSeq_R1.fastq.gz" with a single-end approach. Note: In the command below there is a space before "--" and one before each "/home/user…".

```
bismark --bowtie2 /home/user/DNA/bowtie2-2.3.0/bismark_v0.17.0/REF/
/home/user/SampleSeq_R1.fastq.gz
```

- This aligns the sequence reads to the reference genome and creates a combined alignment/methylation call output in BAM format, and gives a run statistics report.
- Output files: written to /home

    SampleSeq_R1_bismark_bt2.bam

    SampleSeq_R1_bismark_bt2_SE_report.txt

**5.3. Methylation extraction:**

- Extracting methylation information out of the ".bam" file created in step 2. Note: In the command below there is a space before the "-s" , "--", and "/home/user…"

```
bismark_methylation_extractor -s --comprehensive
/home/user/SampleSeq_R1_bismark_bt2.bam
```

- This extracts methylation information from the alignment output of the above NGS file.
- Output files: written to /home

  SampleSeq_R1_bismark_bt2.M-bias.txt

  SampleSeq_R1_bismark_bt2.M-bias_R1.png

  SampleSeq_R1_bismark_bt2_splitting_report.txt

  CHG, CpG and CHH contexts for the SampleSeq_R1_bismark_bt2.txt

**5.4. Generate report:**

- Generating Bismark processing report on read alignment and methylation extraction.

  **bismark2report**

- This gives an overall methylation report on the total number of reads and their alignment and methylation.
- Output file: written to /home

  SampleSeq_R1_bismark_bt2_SE_report.html

**Supplementary Datafile 1.** SampleSeq_R1_bismark_bt2_SE_report.txt

Bismark report for: /home/user/SampleSeq_R1.fastq.gz (version: v0.17.0)
Option '--directional' specified (default mode): alignments to complementary strands (CTOT, CTOB) were ignored (i.e. not performed)
Bismark was run with Bowtie 2 against the bisulfite genome of /home/user/DNA/bowtie2-2.3.0/bismark_v0.17.0/hp21sie1ref/ with the specified options: -q --score-min L,0,-0.2 --ignore-quals

Final Alignment report
======================
Sequences analysed in total: 30135
Number of alignments with a unique best hit from the different alignments: 10571
Mapping efficiency: 35.1%
Sequences with no alignments under any condition: 19564
Sequences did not map uniquely: 0
Sequences which were discarded because genomic sequence could not be extracted: 0
Number of sequences with unique best (first) alignment came from the bowtie output:
CT/CT: 10571 ((converted) top strand)
CT/GA: 0 ((converted) bottom strand)
GA/CT: 0 (complementary to (converted) top strand)
GA/GA: 0 (complementary to (converted) bottom strand)

Number of alignments to (merely theoretical) complementary strands being rejected in total: 0

Final Cytosine Methylation Report

==================================

Total number of C's analyzed: 829618

Total methylated C's in CpG context: 18365

Total methylated C's in CHG context: 647

Total methylated C's in CHH context: 2294

Total methylated C's in Unknown context: 0

Total unmethylated C's in CpG context: 13334

Total unmethylated C's in CHG context: 189351

Total unmethylated C's in CHH context: 605627

Total unmethylated C's in Unknown context: 0

C methylated in CpG context: 57.9%

C methylated in CHG context: 0.3%

C methylated in CHH context: 0.4%

Can't determine percentage of methylated Cs in Unknown context (CN or CHN) if value was 0

Bismark completed in 0d 0h 0m 19s

**Supplementary Datafile 2.** SampleSeq_R1_bismark.bt2.M-bias.txt

CpG context

===========

| position | count methylated | count unmethylated | % methylation | coverage |
|---|---|---|---|---|
| 1 | 0 | 0 | | 0 |
| 2 | 0 | 0 | | 0 |
| 3 | 0 | 0 | | 0 |
| 4 | 0 | 0 | | 0 |
| 5 | 0 | 0 | | 0 |
| 6 | 0 | 0 | | 0 |
| 7 | 0 | 0 | | 0 |
| 8 | 0 | 0 | | 0 |
| 9 | 0 | 0 | | 0 |
| 10 | 0 | 0 | | 0 |
| 11 | 0 | 0 | | 0 |
| 12 | 0 | 0 | | 0 |
| 13 | 0 | 3 | 0.00 | 3 |
| 14 | 0 | 0 | | 0 |
| 15 | 0 | 0 | | 0 |
| 16 | 0 | 0 | | 0 |
| 17 | 0 | 0 | | 0 |
| 18 | 0 | 0 | | 0 |
| 19 | 0 | 0 | | 0 |

| | | | | |
|---|---|---|---|---|
| 20 | 0 | 0 | | 0 |
| 21 | 0 | 0 | | 0 |
| 22 | 0 | 3 | 0.00 | 3 |
| 23 | 0 | 0 | | 0 |
| 24 | 0 | 0 | | 0 |
| 25 | 0 | 13 | 0.00 | 13 |
| 26 | 0 | 0 | | 0 |
| 27 | 1 | 0 | 100.00 | 1 |
| 28 | 0 | 0 | 0 | |
| 29 | 1 | 0 | 100.00 | 1 |
| 30 | 2 | 2 | 50.00 | 4 |
| 31 | 4 | 2 | 66.67 | 6 |
| 32 | 17 | 6 | 73.91 | 23 |
| 33 | 136 | 39 | 77.71 | 175 |
| 34 | 7696 | 2639 | 74.47 | 10335 |
| 35 | 11 | 4 | 73.33 | 15 |
| 36 | 0 | 2 | 0.00 | 2 |
| 37 | 0 | 0 | 0 | |

**Supplementary Datafile 3.** SampleSeq_R1_bismark_bt2.M-bias_R1.png

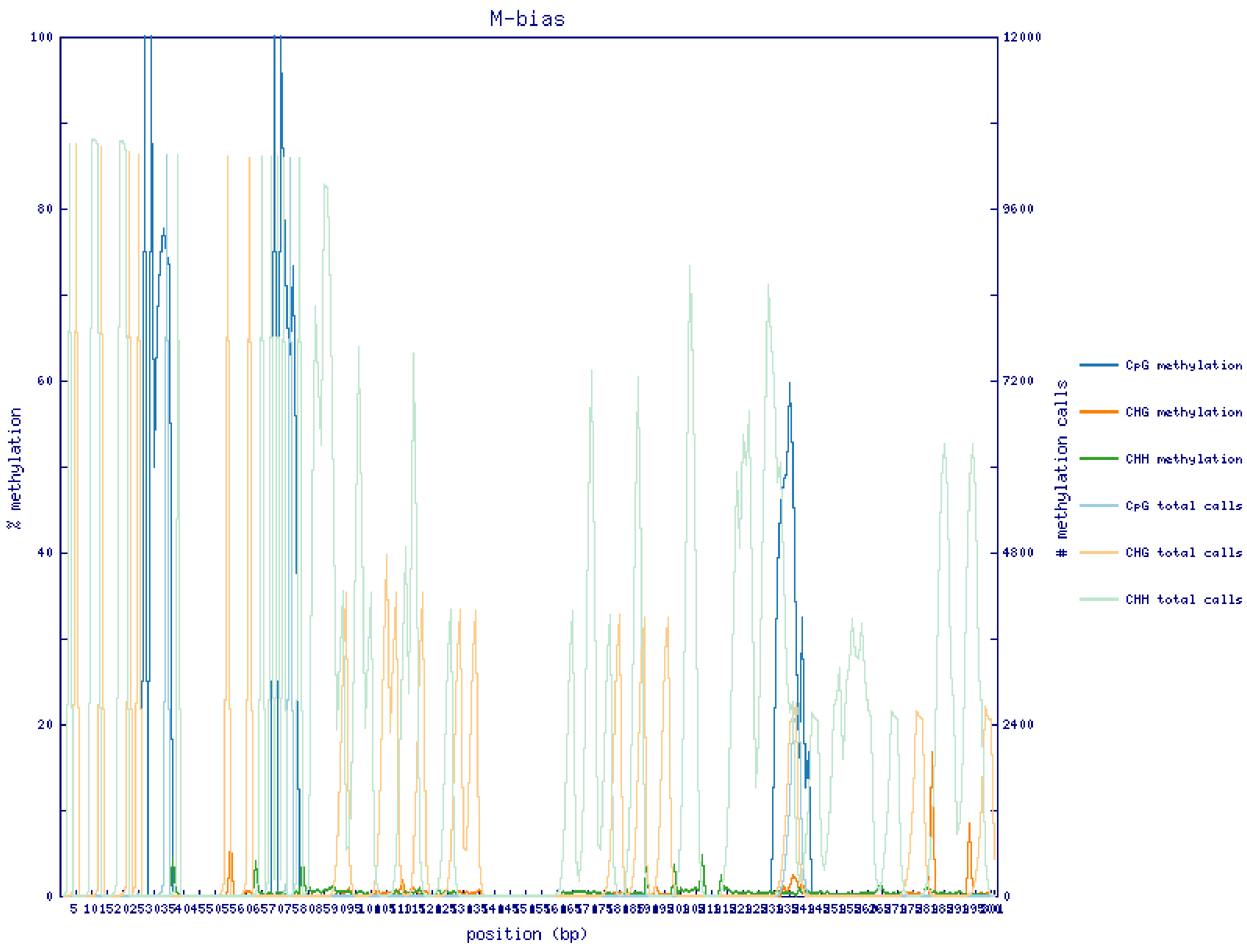

**Supplementary Datafile 4:** CpG_context_SampleSeq_R1_bismark_bt2.txt

Bismark methylation extractor version v0.17.0
M02849:171:000000000-ANHND:1:1101:12957:2144_1:N:0:102 + gi|902576|gb|U24170.1|HSU24170 499 Z
M02849:171:000000000-ANHND:1:1101:12957:2144_1:N:0:102 + gi|902576|gb|U24170.1|HSU24170 539 Z
M02849:171:000000000-ANHND:1:1101:12957:2144_1:N:0:102 + gi|902576|gb|U24170.1|HSU24170 703 Z
M02849:171:000000000-ANHND:1:1101:17290:2154_1:N:0:102 + gi|902576|gb|U24170.1|HSU24170 499 Z
M02849:171:000000000-ANHND:1:1101:17290:2154_1:N:0:102 + gi|902576|gb|U24170.1|HSU24170 539 Z
M02849:171:000000000-ANHND:1:1101:17290:2154_1:N:0:102 - gi|902576|gb|U24170.1|HSU24170 703 z
M02849:171:000000000-ANHND:1:1101:19751:2237_1:N:0:102 - gi|902576|gb|U24170.1|HSU24170 499 z
M02849:171:000000000-ANHND:1:1101:19751:2237_1:N:0:102 + gi|902576|gb|U24170.1|HSU24170 539 Z

**Supplementary Table S1**: **Various combinations of NGS platforms and methylation analysis tools in the recent literature.**

| Platform | Data Analysis | Other tools | Reference |
|---|---|---|---|
| PyroMark Q96 ID | Bismark | Q96 ID software methylKit | Röh *at al* 2017 bioRxiv[1] |
| Ion Torrent PGM | - NextGenMap<br>- BiQ Analyzer HT | Pheatmap | Bhat *et al* 2016[2] |
| Illumina NextSeq | Bismark | NGSUtils, Samtools and Bedtools | Masser *et al* 2016[3] |
| - Illumina MiSeq<br>- PyroMark Q96 MD | BS Seeker | - | Bernstein e*t al* 2015[4] |
| Illumina MiSeq | CLC Genomics Workbench | - | Masser *et al* 2013[5] |

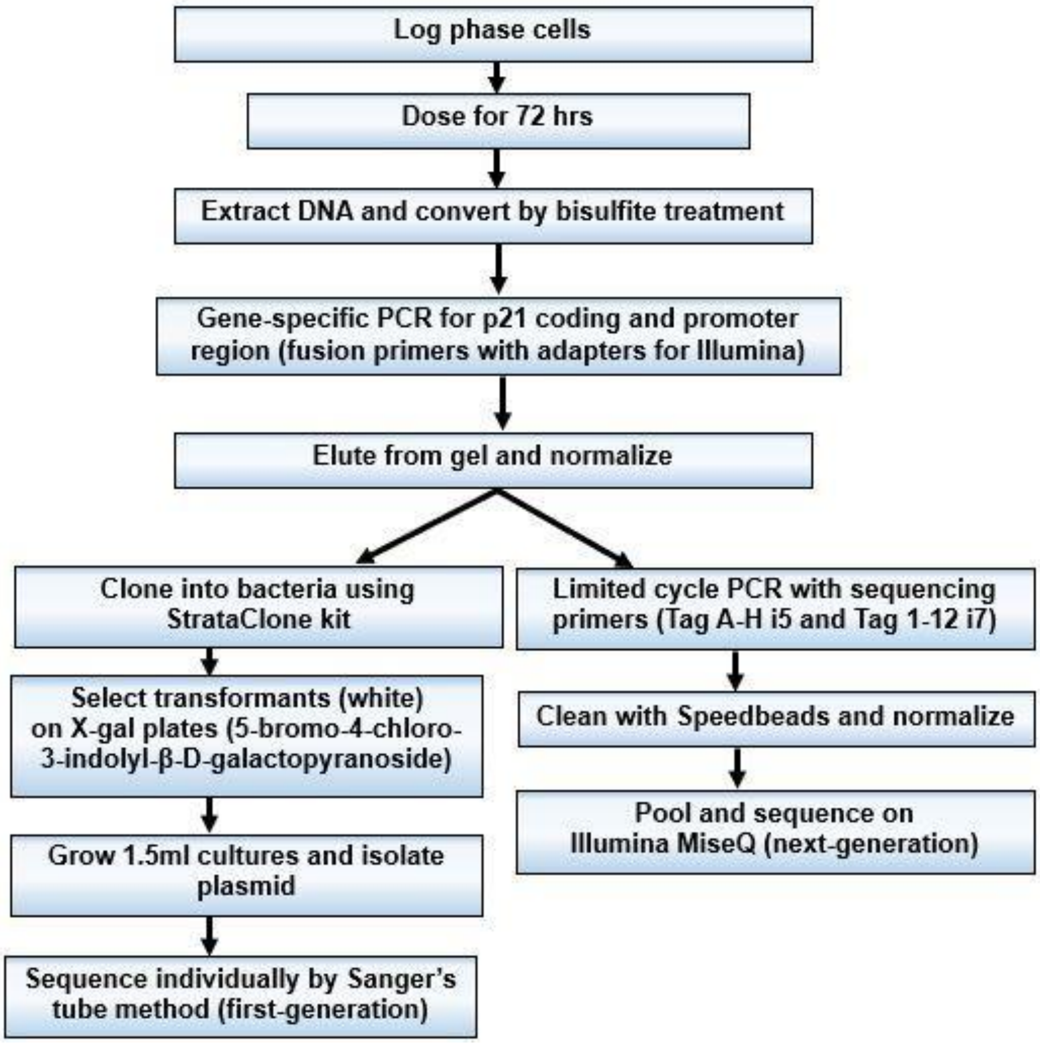


**Supplementary Figure S1**: Schematic for comparing the workflows of first-generation Sanger's and next-generation targeted gene bisulfite sequencing (TGBS) methods.
